# Inhibition of peripheral VEGF signaling rapidly reduces leucocyte obstructions in brain capillaries and increases cortical blood flow in an Alzheimer’s disease mouse model

**DOI:** 10.1101/2021.03.05.433976

**Authors:** Muhammad Ali, Kaja Falkenhain, Brendah N Njiru, Muhammad Murtaza-Ali, Nancy E Ruiz-Uribe, Mohammad Haft-Javaherian, Stall Catchers, Nozomi Nishimura, Chris B. Schaffer, Oliver Bracko

**Affiliations:** Meinig School of Biomedical Engineering, Cornell University, Ithaca, NY, USA, 14853

**Keywords:** Alzheimer’s disease (AD), Amyloid Precursor Protein/ Presenilin 1 (APP/PS1) mouse model, leucocyte, capillary stalling, cerebral blood flow (CBF), Vascular Endothelial Growth Factor (VEGF-A), blood brain barrier (BBB), intraperitoneal, anti-mouseVEGF-A164, endothelial nitric oxide synthase (eNOS), occludin, Evans blue dye

## Abstract

Increased incidence of stalled capillary blood flow caused by adhesion of leucocytes to the brain microvascular endothelium leads to a 17% reduction of cerebral blood flow (CBF) and exacerbates short-term memory loss in multiple mouse models of Alzheimer’s disease (AD). Here, we report that Vascular Endothelial Growth Factor (VEGF) signaling at the luminal side of the brain microvasculature plays an integral role in the capillary stalling phenomenon of the APP/PS1 mouse model. Administration of the anti-mouse VEGF-A164 antibody, an isoform that inhibits blood brain barrier (BBB) hyperpermeability, reduced the number of stalled capillaries within an hour of injection, leading to an immediate increase in average capillary blood flow but not capillary diameter. VEGF-A inhibition also reduced the overall eNOS protein concentrations, increased occludin levels, and decreased the penetration of circulating Evans Blue dye across the BBB into the brain parenchyma, suggesting increased BBB integrity. Capillaries prone to neutrophil adhesion after anti-VEGF-A treatment also had lower occludin concentrations than flowing capillaries. Taken together, our findings demonstrate that VEGF-A signaling in APP/PS1 mice contributes to aberrant eNOS/occludin- associated BBB permeability, increases the incidence of capillary stalls, and leads to reductions in CBF. Reducing leucocyte adhesion by inhibiting luminal VEGF signaling may provide a novel and well-tolerated strategy for improving brain microvascular blood flow in AD patients.

## Introduction

Vascular dysfunction plays a critical role in the pathogenesis of Alzheimer’s disease (AD) and other forms of dementia. Many conditions that drive altered vascular structure and impaired vascular function, including hypertension, obesity, type 2 diabetes, and atherosclerosis, are primary risk factors for AD and other dementias (*1*). In addition, both AD patients and mouse models of AD show cerebral blood flow (CBF) reductions of 10-30%, beginning in the early stages of disease pathogenesis and continuing through disease progression (*2, 3*). Within the AD population, there is also evidence that more severely impaired CBF is associated with poorer cognitive performance (*4*). Even in older humans with no neurodegenerative disorder, lower blood flow in the hippocampus is associated with poorer spatial memory (*5*). Despite decades of data establishing clear links between CBF deficits and greater dementia risk and severity, the underlying mechanisms causing CBF deficits in AD or other dementias are only beginning to be revealed, as are the mechanisms by which vascular risk factors contribute to AD pathogenesis.

We recently found that obstructions in cortical capillary segments occur more frequently in multiple mouse models of AD, as compared to wild type mice. We further found that administering an antibody against the neutrophil-specific cell surface protein lymphocyte antigen 6 complex (Ly6G) led to a ~60% decrease in the incidence of non-flowing capillaries and a ~20% increase in CBF, all within 10 minutes. This CBF increase was correlated with striking improvements in performance on spatial and short-term memory tasks in the AD mice that were apparent at three hours after anti-Ly6G administration (*6*). This work has suggested that interfering with neutrophils adhering in capillary segments could be a novel therapeutic approach for AD, making efforts to elucidate the molecular signaling that links AD pathogenesis to increased incidence of capillary stalling critical.

Vascular endothelial growth factor (VEGF-A) is involved in signaling pathways that drive a broad variety of physiological processes including angiogenesis (*7*), vascularization (*8*), lymphangiogenesis (*9*), growth tip guidance (*10*), vascular dysfunction (*8*), response to fluid shear stress (*11*), cellular junction integrity (*8*), neurogenesis (*10*), and neuroprotection (*7*). With such a complex set of downstream impacts, it is not surprising that alterations of VEGF-A signaling have been found to have a wide range of effects in the CNS and in the periphery. VEGF-A is secreted into the blood from many tissues and several transport routes such as lymphatic drainage, microvascular permeability, internalization, and plasma clearance. The source and route of VEGF-A transportation is crucial for its function. For example, VEGF-A levels in the blood regulate blood brain barrier integrity(*12–14*). In mouse models of diabetic retinopathy, peripherally inhibiting VEGF-A signaling reduced the incidence of retinal capillary obstructions caused by adhered leucocytes (*15*). Additionally, peripheral inhibition of vascular endothelial growth factor receptor-2 (VEGF-2R) lead to improved clearance of microsphere-induced cortical capillary obstructions and a reduction in the pruning of such obstructed capillaries (*16, 17*). These findings suggest that peripheral VEGF-A signaling plays a critical role in microvascular obstructions and that inhibiting VEGF-A signaling may improve microvascular flow in the central nervous system (CNS).

Differing changes in and impacts of VEGF-A signaling have been observed in AD, depending largely on whether the focus is on VEGF-A signaling in the periphery or in the brain, itself. Several reports have shown increased VEGF-A levels in the blood plasma of AD patients (*18–21*), while only a few found the opposite (*22*). Expression of VEGF-A was also found to be increased in capillaries and blood vessels from AD patients (*23, 24*). This increased VEGF-A signaling could contribute to capillary obstructions, as in mouse models of diabetic retinopathy (*15*). On the other hand, VEGF-A signaling plays a supportive role for cells within the CNS, and increased VEGF-A signaling in the brain has been shown to have beneficial effects in AD mouse models. Specifically, VEGF-A-releasing nanoparticles injected into the brain of APP/PS1 mice improved behavioral deficits and reduced neuronal loss (*25*). Here, we hypothesized that inhibiting peripheral VEGF-A signaling may reduce the incidence of non-flowing capillaries in the APP/PS1 mouse models of AD, thereby increasing CBF.

Using in vivo two-photon excited fluorescence (2PEF) microscopy, we examined the impact of peripheral VEGF-A inhibition on the incidence of non-flowing capillaries in the brain microvasculature of the APP/PS1 mouse model of AD. We used an antibody against the murine VEGF-A165 protein (which we refer to as “anti-VEGF-A” in this manuscript) that is too large to cross the BBB in significant amounts (*26–28*). We quantified the incidence of non-flowing capillaries as well as blood flow speeds in flowing capillaries before and after administration of antibodies against VEGF-A. To unearth potential mechanisms by which VEGF-A regulates capillary obstructions and CBF in the CNS, we examined the expression of proteins associated with the BBB and their function in APP/PS1 mice and controls, as well as from human tissue samples from AD patients.

## Methods

### Animals

All animal procedures were approved by the Cornell Institutional Animal Care and Use Committee (protocol number 2015-0029) and were conducted under the oversight of the Cornell Center for Animal Resources and Education. We used the APP/PS1 transgenic mouse model of AD (B6.Cg-Tg (APPswe, PSEN1dE9) 85Dbo/J; MMRRC_034832-JAX, The Jackson Laboratory), with wild type (WT) littermates as controls.

### Experimental Cohorts

The experiments were conducted in three cohorts. The first cohort of mice (10-14 months) was divided into four experimental groups: AD mice given the anti-VEGF-A treatment (n=5; 3 male, 2 female), AD mice given a placebo saline injection (n=4; 2 male, 2 female), WT mice given the anti-VEGF-A treatment (n=6; 3 male, 3 female), and WT mice given a placebo saline injection (n=6; 3 male, 3 female) (Figure 2A). Each anti-VEGF-A injection consisted of 200 μL of 32.5 pM concentration of antibody against mouseVEGF-A164 protein (YU1418071, R&D Systems, Inc.) in saline, and was administered intraperitoneally. Control injections consisted of 200 μL of saline. Mice first received cranial windows for imaging (see below) and following a 2-3-week recovery period, anti-VEGF-A injections were administered every other day over a two-week span. Mice were imaged at the one-week mark, which was after the first three injections, and again at the two-week mark, after the last three injections. After the second imaging session, these mice were sacrificed and their brains were removed for ELISA assays and immunofluorescence.

The second cohort of mice (12-15 months) was divided into two experimental groups: AD mice (n=9; 5 male, 4 female) and WT mice (n=4; 2 male, 2 female) (Figure 5A). After receiving a cranial window and being given 2-3 weeks to recover, these animals also received anti-VEGF-A injections every two days for one week, with the same dosage and route of administration as above. With this cohort, mice were first imaged at baseline, 1 hr. after the first injection, and finally at one week after a total of three injections. After the final imaging sessions, these animals were sacrificed and their brains were perfused for qualitative Evan’s Blue leakage analysis (see below).

The third cohort of mice (9-10 months) was divided into three experimental groups: AD mice given the anti-VEGF-A treatment (n=3; 2 male, 1 female), AD mice given a placebo saline injection (n=3; 2 male, 1 female), WT mice given the anti-VEGF-A treatment (n=3; 2 male, 1 female), and WT mice given the placebo saline injection (n=3; 2 male, 1 female) (Figure 5B). These animals were sacrificed one hour after treatment and their brains were perfused for quantitative Evan’s Blue leakage analysis.

### Surgical Procedure

The animals were initially anesthetized with 3% isoflurane and maintained on 1.5% isoflurane in oxygen. They were injected subcutaneously with atropine (54925-063-10, Med-Pharmex), dexamethasone (07-808-8194, Phoenix Pharm) and ketoprofen (Zoetis). The atropine was administered at 0.005 mg per 100-g mouse weight and was used to prevent fluid buildup in the lungs. The dexamethasone was administered at 0.025 mg per 100-g mouse weight and was used to reduce surgery-induced inflammation. The ketoprofen was administered at 0.5 mg per 100-g mouse weight and was used to reduce surgery-induced inflammation and to provide post-surgical analgesia.

The top of the head was shaved using clippers (Oster). The exposed area was washed three times, alternating between a 70% ethanol and an iodine solution (AgriLabs). The mice were then placed on a feedback-control heating blanket (40-90-8D DC, FHC) to maintain body temperature at 37°C. The head was fixed on a custom-built stereotactic surgery frame using ear bars and a bite bar. Bupivacaine (0.1 ml of a 0.125% solution in saline; Hospira) was injected subcutaneously in the scalp to provide a local nerve block. The skin was opened and a 6 – 7-mm diameter craniotomy was performed on the top of the skull, centered on the midline and about halfway between the lambda and bregma points. This was done with a high-speed drill (HP4-917-21, Fordom), using 0.9-mm diameter drill bits at the top of the skull and switching to 0.7-mm and, finally, 0.5-mm diameter bits when nearly through the skull (Fine Science Tools). The circumscribed bone flap was removed, the brain covered with saline, and the opening covered with an 8-mm diameter glass coverslip (11986309, Thermo Scientific), which was glued to the remaining skull using cyanoacrylate adhesive (Loctite) and dental cement (Co-Oral-Ite Dental).

For two days after surgery, the animals were treated with dexamethasone at 0.025 mg per 100-g animal weight and ketoprofen at 0.5 mg per 100-g animal weight, daily. The mice were given at least two weeks to recover from this surgery. If the clarity of the optical window was deemed too poor for high quality 2PEF imaging after the two weeks, the mice were sacrificed and discontinued from the study (6 out of 34 mice were removed due to poor cranial windows).

### *In Vivo* Two-Photon Microscopy

The animals were initially anesthetized with 3% isoflurane and maintained on 1.5% isoflurane. They were then placed on a custom-built stereotactic surgery frame, injected with atropine as described above, and were placed above a feedback-control heating blanket maintaining body temperature at 37°C. For fluorescent labeling of the microvasculature, 70-kDa Texas Red dextran (40 μL of a 2.5% w/v solution in saline; Thermo Fischer Scientific) was retro-orbitally injected before imaging. Some animals were also retro-orbitally injected with a second syringe containing Rhodamine 6G (0.1 ml of a 1-mg ml^−1^ solution in saline; Acros Organics) to distinguish circulating leucocytes and platelets from red blood cells (RBCs) *via* mitochondrial labeling. Imaging was conducted using 830-nm, 75-fs pulses from a Ti:Sapphire laser oscillator (Vision S, Coherent). The laser beam was raster scanned using galvanometric scan mirrors at one frame per second. Microvasculature was imaged using a 20X water-immersion objective lens for high-resolution imaging (numerical aperture of 1.0, Carl Zeiss; or numerical aperture of 0.95, Olympus), or a 4X objective (numerical aperture of-0.28, Olympus) for broader mapping of the vascular network on the surface of the cortex. The emitted fluorescence was detected using detection optics that were ray traced to efficiently deliver the divergent cone of light emerging from the back aperture of the objective to the ~5-mm diameter active area of high quantum efficiency GaAsP photomultiplier tubes (H6527, Hamamatsu).

Fluorescence signaling from the Texas Red dye was detected through a 641/75 nm (center wavelength/bandwidth) filter and the fluorescence from Rhodamine 6G was detected through a 550/49 nm filter, with the light separated by a long-pass dichroic with a cutoff wavelength of 605 nm. ScanImage software was used to control the laser scanning and data acquisition(*29*). Three-dimensional images of the brain microvasculature (Figure 1A) and line scan measurements of RBC speed in capillaries (Figure 2C) were acquired using this system. Stacks of images were spaced 1 μm axially and were taken to a depth of 200 – 400 μm. From within this imaging volume, 6-10 capillary segments, all with a diameter of less than 10 μm, were randomly selected for line scan-based measurement of flow speed and for small image stacks to quantify vessel diameter.

**Figure 1.**
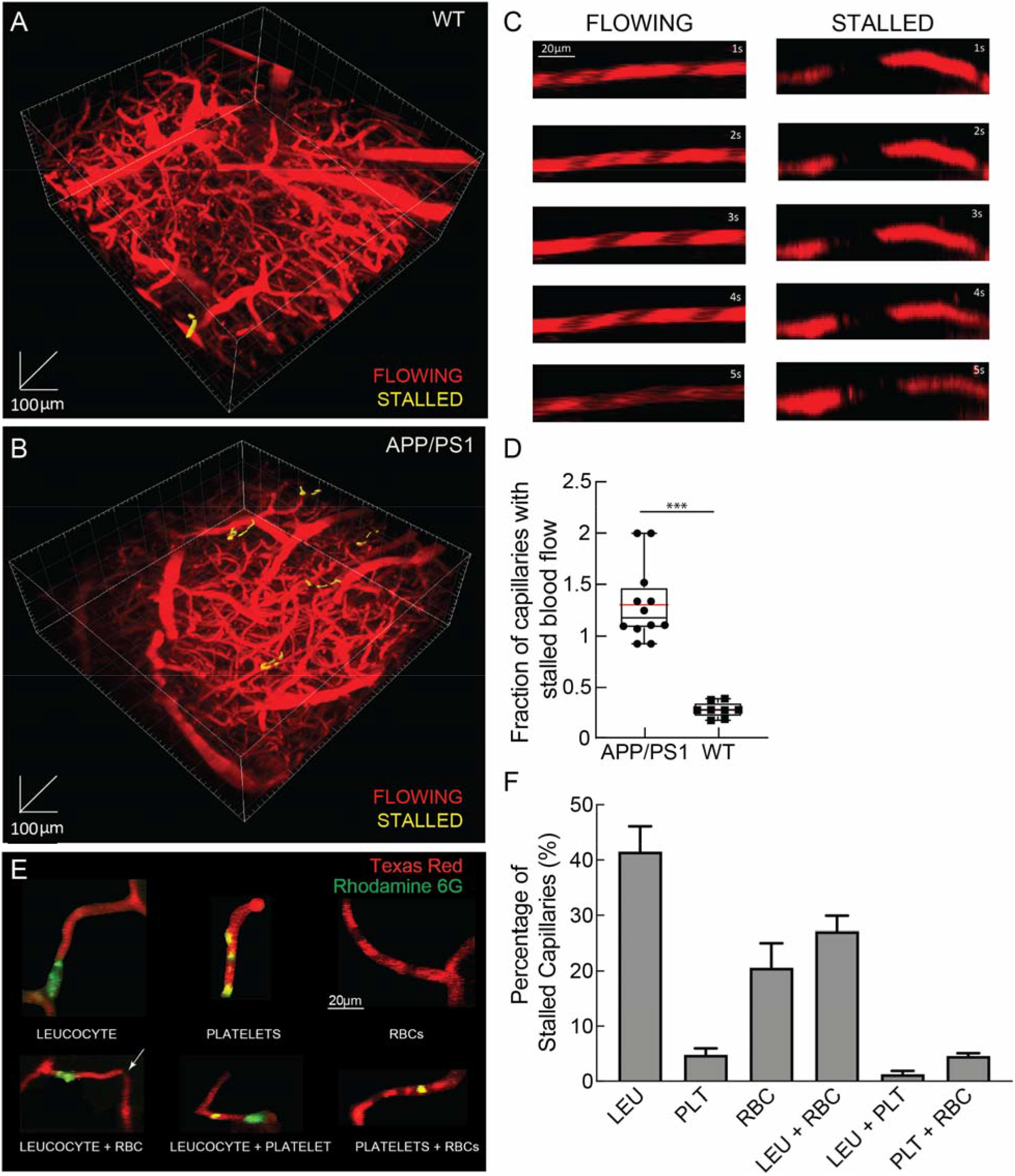
2PEF imaging of mouse cortical vasculature showed a larger fraction of capillaries with stalled blood flow in APP/PS1 mice. Rendering of a 2PEF image stack of the cortical vasculature (red; Texas Red dextran) from a WT mouse (**A**), with a single stalled capillary indicated in yellow, and of an APP/PS1 mouse (**B**), with five stalled capillaries. (**C**) Individual capillaries throughout the image stack were characterized as flowing or stalled based on the movement of unlabeled (black) red blood cells within the fluorescently-labeled blood plasma (red). (**D**) Fraction of capillaries with stalled blood flow (APP/PS1 (n=12) and WT (n=8) mice, ~ 23,000 capillaries; two-tailed Mann-Whitney test, p *=* 0.001; boxplot: whiskers extend 1.5x the difference between the 25^th^ and 75^th^ percentile of the data, the red horizontal line represents the median, the black line represents the mean). (**E**) Z projection of image stacks through stalled capillaries that contain a leucocyte (LEU, upper left), platelet aggregates (PLT, upper center), RBCs (upper right), LEU and RBCs (lower left), LEU and PLT (lower center), and PLT and RBCs (lower right), distinguished by fluorescent labels (red: Texas Red labeled blood plasma; green: Rhodamine 6G labeled LEU and PLT). (**F)** Percentage of capillary stalls in APP/PS1 mice that contained only LEU, only PLT, only RBCs, both LEU and RBCs, both LEU and PLT, and both PLT and RBCs (n=4, 98 capillaries).

**Figure 2.**
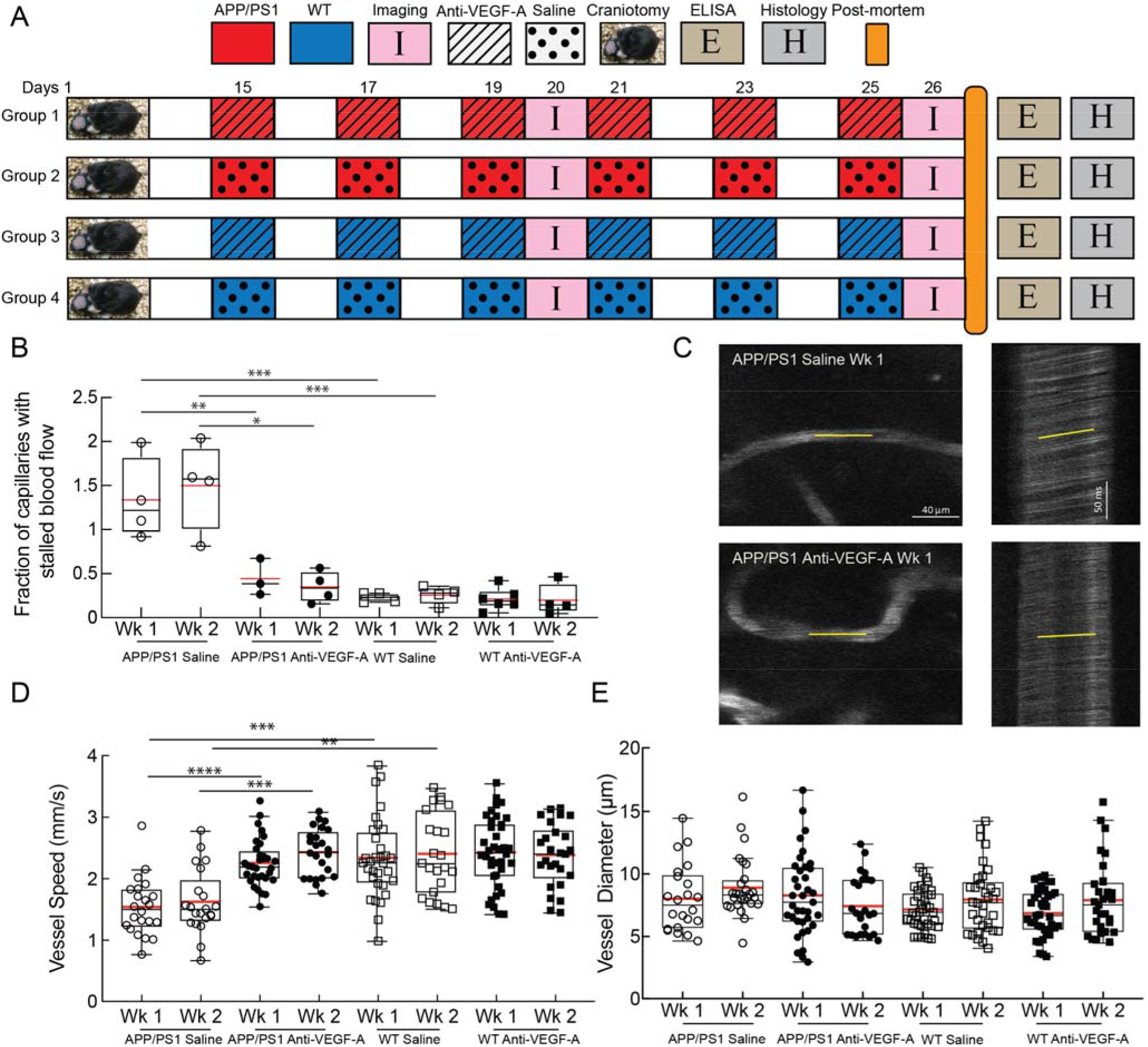
Anti-VEGF-A antibody treatment reduced the incidence of capillary stalling increased cerebral blood flow speed in APP/PS1 mice. (**A**) **S**chematic of experimental timeline. After craniotomies were performed and the mice recovered for two weeks, the mice were divided into four groups and treated, as follows, every other day for two weeks: APP/PS1 mice injected with saline (n=4; APP/PS1-saline), APP/PS1 mice treated with anti-VEGF-A (n=5; APP/PS1-anti-VEGF-A), WT mice injected with saline (n=6; WT-saline), and WT mice treated with anti-VEGF-A (n=6; WT-anti-VEGF-A). These mice were imaged twice, on the sixth and twelfth day after the first injection. Their brains were harvested for post-mortem assays after the second imaging session. (**B**) Box plot of the fraction of capillaries with stalled blood flow after 1 week and 2 weeks of anti-VEGF-A or saline treatment in APP/PS1 and WT mice ((APP/PS1-anti-VEGF-A: 2 mice excluded at week 1 timepoint due to motion artifact, 1 mouse lost cranial window before week 2 timepoint; WT-saline: 1 mouse at week 1 excluded due to motion artifact, 1 mouse lost cranial window before week 2; WT-anti-VEGF-A: 1 mouse lost cranial window before week 2 imaging session); ~ 48,000 capillaries; one way analysis of variance (ANOVA) with Holm-Šídák post hoc multiple comparison correction to compare across multiple groups: week 1 APP/PS1 saline vs. week 1 APP/PS1-anti-VEGF-A p value = 0.004, week 2 saline vs. week 2 anti-VEGF-A APP/PS1 p value = 0.001, week 1 saline APP/PS1 vs. week 1 saline WT p value = 0.001, week 2 APP/PS1-saline vs. week 2 WT-saline p value = 0.001, week 1 APP/PS1-anti-VEGF-A vs. week 1 APP/PS1-saline p value = 0.01; each data point in the graph represents the fraction of stalled capillaries in 4-6 2PEF stacks for each mouse). (**C**) Images (left) and line-scans (right) from representative vessels from a saline (top) and anti-VEGF-A (bottom) injected APP/PS1 mouse. (**D**) RBC flow speed and (**E**) vessel diameter in cortical capillaries after 1 week and 2 weeks of anti-VEGF-A or saline treatment in APP/PS1 and WT mice (week 1 APP/PS1-saline: n=4, 22 vessels, week 2 APP/PS1-saline: n=4, 21 vessels, week 1 APP/PS1-anti-VEGF-A: n=5, 32 vessels, week 2 APP/PS1-anti-VEGF-A: n=4, 24 vessels (1 mouse lost cranial window after week 1 imaging session), week 1 WT-saline: n=6, 32 vessels, week 2 WT-saline: n=5, 23 vessels (1 mouse lost cranial window after week 1 imaging session), week 1 WT-anti-VEGF-A: n=6, 38 vessels, week 2 WT-anti-VEGF-A: n=5, 26 vessels (1 mouse lost cranial window after week 1 imaging session); one way ANOVA with Tukey’s post hoc multiple comparison correction to compare vessel speed across groups: week 1 saline APP/PS1 vs. week 1 anti-VEGF-A APP/PS1 p value < 0.0001, week 2 saline vs. week 2 APP/PS1-anti-VEGF-A p value = 0.0010, week 1 WT-saline vs. week 1 WT-anti-VEGF-A p value > 0.99, week 2 saline WT vs. week 2 WT-anti-VEGF-A p value > 0.99, week 1 APP/PS1-saline vs. week 1 WT-saline p value = 0.0007, week 2 APP/PS1-saline vs. week 2 WT-saline p value = 0.0022, week 1 APP/PS1-anti-VEGF-A vs. week 1 WT-saline p value > 0.99, week 2 APP/PS-anti-VEGF-A 1 vs. week 2 WT-saline p value > 0.99; each point in the graph represents one of the 4-8 capillaries measured in each mouse at each imaging session). In all graphs the boxplot whiskers extend 1.5x the difference between the 25^th^ and 75^th^ percentile of the data, the red horizontal line represents the median, and the black line represents the mean.

### Analysis of capillary speed and diameter

We identified capillary segments that were largely parallel to the cortical surface. We took a small image stack that was later used to determine the diameter of the vessel using the plot profile option on ImageJ. Speeds of RBC flow in these capillaries were determined by repeatedly scanning a line along the central axis of the vessel. Unlabeled RBCs showed up as dark patches inside the vessel lumen, and the movement of these unlabeled RBCs was tracked over successive line scans by constructing a space-time plot, where the x-axis represented distance along the vessel axis and the y-axis represented time. Diagonal stripes on this plot were formed by the moving RBCs and had a slope that was inversely proportional to the RBC speed, which was calculated using a Radon transform-based algorithm that has been previously described by (*30*).

### Crowd-sourced quantification and analysis of capillary stalling

To determine the incidence of non-flowing capillaries within the microvasculature we used a purpose-built citizen science data analysis platform, StallCatchers. We previously detailed and validated this methodology (Bracko et al., 2019). Briefly, we used the convolutional neural network DeepVess to segment 2PEF stacks into voxels defined to be within or outside of the vasculature (*31*). Vessel centerlines were then determined from this segmented image using standard dilation and thinning operations (*32*), and individual vessel segments were identified. Vessels with diameters greater than 10 μm were excluded in order to restrict our analysis of stalling behaviors to capillaries. Identified capillary segments were outlined and a cropped image stack centered on each outlined capillary was uploaded to www.StallCatchers.com. Volunteers scored each of the ~60,000 capillaries examined in this study as either flowing (0) or stalled (1). Since the Texas Red dextran labels the blood plasma, blood cells were seen as dark patches in the microvasculature lumen. StallCatchers users would determine whether they saw these dark patches move along the vessel axis across the multiple frames in which the outlined capillary was present (minimum of ~5). Multiple volunteers scored each capillary as flowing or stalled. Each volunteer had a unique sensitivity score determined by their performance on intermixed capillaries that had already been scored as stalled or flowing by lab members. A weighted “crowd-confidence” score that took each volunteer’s sensitivity into account, ranging from 0 (likely flowing) to 1 (likely stalled) was then determined for each capillary. Capillaries with scores above 0.5 (n=900) were validated by lab members who were blinded to the genotype and treatment-status. Capillaries with crowd confidence scores of 0.9-1 were found to be stalled 95% of the time by lab members, whereas with confidence scores between 0.5 and 0.6, only 1.5% of the capillaries were determined to be stalled by lab members. This citizen science scoring of capillary flow reduced the number of capillaries directly examined by lab members by a factor of 50. Finally, stall rates were calculated as the number of stalled capillaries divided by the number of total capillaries across all 2PEF images taken for each mouse at each time point. All imaging stacks were manually checked for wrong connection after segmentation.

Upon identifying capillary stalls, in certain animals the Rhodamine 6G channel was used to determine the fraction of stalled capillaries that contained only leucocytes, only platelets, only RBCs, both leucocytes and RBCs, or both platelets and RBCs, using the distinguishing criteria previously described by (*33*). All image analysis was conducted using ImageJ software (NIH).

### Analysis of vascular density

We calculated the averaged density of the capillary network using the DeepVess (*31*) for vascular segmentation. We reported the capillary density as the fraction of image voxels determined to be inside vessels divided by the total number of image voxels. Image stacks were first manually masked to exclude large surface vessels from this analysis.

### ELISA assay

After two weeks of treatment, the mice from the first cohort were sacrificed by a retro-orbital injection of pentobarbital at 5 mg per 100 g mouse weight. Brains were extracted and cut along the centerline. One half was preserved in 4% paraformaldehyde (PFA) in 1X phosphate buffered saline (PBS) for 24 hrs, and then stored in 30% sucrose solution for histology(*34*).

The other half was weighed and homogenized using a Dounce homogenizer (8343-15, ACE GLASS, INC.) in 1 ml PBS with complete protease inhibitor (Roche Applied Science) and 1 mM of the serine protease inhibitor 4-(2-aminoethyl) benzenesulfonyl fluoride hydrochloride (ThermoFisher Scientific). The samples were sonicated and centrifuged at 14,000g for 30 min at 4°C. The PBS-soluble supernatant was separated and stored at −80°C. The remaining pellet was dissolved in 0.5 ml 70% formic acid before being sonicated and centrifuged at 14,000g for 30 min at 4°C. The supernatant was separated and neutralized in 1M Tris buffer at a pH of 9.0. Protein concentration was measured using the Pierce BCA Protein Assay (ThermoFisher Scientific). The brain extracts were diluted to equal protein concentration and were analyzed by sandwich ELISA for VEGF-A, endothelial nitric oxide synthase (eNOS), amyloid-beta (Aβ) 40, Aβ42, and Aβ aggregates using commercial ELISA kits and following the manufacturer’s protocols (VEGF-A: ab209882, Abcam; eNOS: ab230938, Abcam; Aβ40: KHB3481, and Aβ42: KHB3441, ThermoFisher Scientific).

Tissue from brain lysates and plasma samples was collected as previously described(*34*). Brains were weighed and homogenized using a Dounce homogenizer (8343-15, ACE GLASS, INC.) in 2 ml PBS with complete protease inhibitor (Roche Applied Science), PhosSTOP (Roche Applied Science), and 1 mM of the serine protease inhibitor 4-(2-aminoethyl) benzenesulfonyl fluoride hydrochloride (ThermoFisher Scientific). The samples were further ruptured using a syringe and centrifuged at 14,000g for 30 min at 4°C. The PBS-soluble supernatant was separated and stored at −80°C. After the measuring the concentrations ELISAs were performed following the manufacture’s protocol (VEGF1R (MVR100) and VEGF2R (MVR200B), R&D Systems). Peripheral blood drawn using cardiac puncture and centrifuged for 10 min at 3,000 rpm (1500 × g) and the plasma was transferred to a new tube. The albumin was removed using the Pierce™ Albumin Depletion Kit (ThermoFisher Scientific). After the measuring the concentrations ELISA’s were performed following the manufactures protocol (sVEGF1R (MBS4503308) and sVEGF2R (MBS776967), MyBiosource). Data was collected through a Synergy HT plate reader (Biotek) and analyzed via Gen5 software (BioTek).

### Immunohistochemistry and confocal microscopy

The two-week treated half-brains preserved for immunofluorescence were placed in optimal cutting temperature (OCT) compound (Fisher Scientific) and sliced into 30 μm sections. The sections were mounted and washed in PBS. They were incubated in blocking solution, 250 μl of donkey serum and 5 μL of 100% triton in 5 ml of PBS, for 1 hr., and then overnight at 4°C in a 1:100 primary antibody solution of the anti-GLUT-1 capillary marker antibody (ab15309, Abcam) and the BBB integrity protein occludin anti-Occludin antibody (ab31721, Abcam) in blocking solution. The following day, the samples were washed in PBS and incubated for 1.5 hr in a 1:300 secondary antibody solution of a 488 nm anti-mouse antibody (ThermoFisher Scientific) and a 594 nm anti-rabbit antibody (ThermoFisher Scientific). The sections were washed 3x in PBS and allowed to thoroughly dry and coverslips were added onto the slides.

The slides were imaged using confocal microscopy on three-channels. Z-stacks were taken from the pre-frontal cortex, somatosensory cortex, parietal lobe, and hippocampus up to 30 μm deep. The same gain and excitation limits were used across all samples. For broad analysis of the vasculature, each stack of images was converted to a single Z project. The Renyi entropy filter on ImageJ was applied, and individual particles were outlined. Integrated density was calculated for each channel (Supplementary Figure 1). Results were reported as “Integrated density of occludin/ Integrated density of GLUT-1.” Analysis of specific vessels entailed outlining entire capillaries, from one end to the other. ImageJ’s mean gray value function was applied to this outline, specifically on the occludin channel. These results were reported as “Mean occludin intensity throughout capillary length.”

### Human staining

Post-mortem human cortical tissue from males and females were obtained through the NIH NeuroBioBank. Frozen cortical brain samples were fixed in formalin or were received in formalin from the brain bank. 30 μm thick sections were sectioned in a cryostat and stored in cryoprotection solution until further processing. Cortical brain sections were rinsed for 5 min with 1X PBS and incubated with a 0.3% solution of Sudan Black B in 70% ethanol for 30 min to reduce lipofuscin autofluorescence. Slices were then blocked in 3% goat serum for 30 min and incubated with primary antibody overnight at 4°C. On the next day, slices were rinsed and incubated with secondary antibody (goat anti-rabbit Alexa Fluor 594) at room temperature for 2 hours. Rabbit anti-occludin antibody (abcam, ab222691and Lectin (Thermofisher, CAT # L32482)) was used for blood vessel labeling at a 1:250 dilution in blocking solution. Slides were allowed to dry, mounted with Prolong Gold antifade mountant (P36394) imaged with a Zeiss 710 confocal microscope. We used the Renyi entropy thresholding method to define labeled pixels and calculated the integrated density for each fluorescence channel. Eight fields of view across four slices were averaged for each sample. Results are reported as “Integrated density of occludin / Integrated density of lectin”. To determine statistical significance, a two-tailed unpaired t-test was performed.

### Evan’s Blue leakage analysis

After the conclusion of one week of treatment, the mice in the second cohort were retro-orbitally injected with Evan’s Blue dye (EB), 2mL/kg of a 2% solution, to track BBB permeability. One hour after EB injection, they were sacrificed by injection of pentobarbital at 5 mg per 100 g. They were transcardially perfused with 1X PBS followed by 4% PFA. Brains were kept in 4% PFA for 24 hours and then transferred to 30% sucrose solution. Brain tissue was cut along the sagittal centerline. Pictures of the half-brains were taken for qualitative analysis.

One hour after an injection of anti-VEGF-A, the mice in cohort 3 were injected with EB, 2 mL/kg of a 2% solution, to track BBB permeability. These mice were similarly sacrificed, but only perfused with 1xPBS. After beheading the mice, the cortex and hippocampus were separated and weighed. The brain sections were then homogenized in a 3:1 PBS: trichloroacetic acid (TCA) (Sigma-Aldrich) solution to precipitate out macromolecular compounds. Samples were cooled overnight at 4°C and then centrifuged at 1000g for 30 min. EB absorbance was quantitatively measured at 620 nm and then divided by the mass of the brain tissue. This methodology has been previously described(*35*).

### Statistics

The Mann-Whitney U-test (two groups) and analysis of variance (ANOVA) test with Dunn’s nonparametric or Tukey’s parametric multiple comparison correction (three or more groups) was used to determine statistical differences between groups of data. For longitudinal studies, the ratio paired t-test (two groups) or ANOVA repeated measures option with equal variability of differences (three or more groups) was used. A standard indicator of statistical significance was used throughout the figures (^*^*p-value*<0.05, ^**^*p-value*<0.01, ^***^*p-value*<0.001, ^****^*p-value*<0.0001). If the derived p-value was less than 0.05 the difference was considered significant. If the p-value was other than such the data sets were assumed to have no difference. Individual data points are presented in graphical form, with a red line representing the mean. Boxplot whiskers extend 1.5x the difference between the 25^th^ and 75^th^ percentile. Longitudinal data are presented in graphs where a solid black line tracks the same mouse over time. Graphpad’s Prism 8 software was used for all statistical analyses and graph design.

## Results

### Anti-VEGF-A treated APP/PS1 mice had a reduced incidence of capillary stalls and increased cortical capillary flow speeds

We used *in vivo* 2PEF microscopy to image mouse cortical microvasculature and observed that only 0.3% of capillaries in WT mice had obstructed flow (Figure 1A), whereas 1.5% of capillaries in APP/PS1 mice were stalled (Figure 1B) (Mann Whitney p-value < 0.001) (Figure 1D). A majority of the capillaries without flow had leucocytes plugging the vessel lumen (~70%), either with or without one or more RBCs present (Figure 1E, 1F). These observations are consistent with our previous findings in APP/PS1 mice, where we demonstrated that 1.8% of capillaries were stalled, leading to reduced blood flow in up- and down-stream vessels and causing a 17% reduction in cortical perfusion, as measured by ASL-MRI (*6*).

We then inhibited VEGF-A signaling for two weeks by administering anti-VEGF-A antibodies (YU1418071, R&D Systems, Inc.; intraperitoneal every other day), and measured the incidence of capillary stalling, capillary blood flow speeds after one and two weeks of treatment (Figure 2A). APP/PS1 mice treated with anti-VEGF-A for one and two weeks showed reduced incidence of capillary stalls, as compared to saline-treated control (one week: 1.34% vs. 0.44%; two weeks: 1.49% vs. 0.38%; one-way ANOVA with Holm-Šídák post hoc multiple comparison correction; p-value = 0.05 and 0.01, respectively) (Figure 2B). In contrast, WT mice had low levels of capillary stalls (0.3%) that were not altered by either anti-VEGF-A or saline treatment. Strikingly, anti-VEGF-A treated APP/PS1 mice had capillary stall incidence that was on par with that of WT mice. The anti-VEGF-A treated APP/PS1 mice also had, on average, 1.5 times faster capillary flow speeds, as compared to saline-treated APP/PS1 mice, after one and two weeks of treatment (one week: 2.3 mm/s vs. 1.5 mm/s; two weeks: 2.4 mm/s vs. 1.8 mm/s; one-way ANOVA with Tukey’s *post hoc* multiple comparison correction; p-value < 0.001 and 0.01, respectively) (Figure 2C and D). Strikingly, the anti-VEGF-A treatment completely eliminated the deficit in capillary flow speeds APP/PS1 mice exhibited relative to WT mice. The anti-VEGF-A treatment did not affect capillary flow speeds in WT mice (Figure 2D) or capillary diameter in either APP/PS1 or WT mice (Figure 2E).

### Anti-VEGF-A treated APP/PS1 mice had reduced eNOS and increased occludin levels

After in vivo imaging, the mice were euthanized and their brain tissue was processed for measuring protein concentration and histology. The anti-VEGF-A treatment lowered the concentration of VEGF-A in whole brain extracts from APP/PS1 mice by 70%, as measured by ELISA and compared to saline-treated APP/PS1 mice, which brought the VEGF-A concentration in line with that from saline-treated WT mice (Figure 3A). Endothelial nitric oxide synthase (eNOS) was reduced by 70% with anti-VEGF-A treatment in APP/PS1 mice, as measured by ELISA and compared to saline treatment (Figure 3B). A similar fractional reduction in eNOS was observed in WT mice treated with anti-VEGF-A. Saline-treated APP/PS1 mice had ~1.5 times higher eNOS concentration than saline-treated WT mice. In histological sections, we quantified the expression of occludin in capillaries from the cortex and hippocampus (Figure 3C). Anti-VEGF-A treatment of APP/PS1 mice increased occludin concentration by 2.5 and 2.0 times in the cortex and hippocampus, respectively, as compared to saline treatment, bringing occludin concentration to the same level as in WT mice (Figure 3D and E). We also measured protein levels of the VEGF 1 and 2 receptors from brain and plasma samples. There was an overall lower concentration of VEGF1R receptors in brain lysates in AD as compared to WT mice, but no notable change with anti-VEGF treatment after one week of treatment (Supplementary Figure 2A). In contrast, there were higher concentrations of VEGF2R levels in APP/PS1 mice compared to WT mice and a trend toward reduced VEGF2R protein levels in APP/PS1 mice after anti-VEGF treatment (Supplementary Figure 2B). Next, we tested for the soluble forms of these two receptors from plasma samples but did not detect any differences between APP/PS1 mice and WT mice or between anti-VEGF treated mice and saline-injected (Supplementary Figure 2C and D). In addition, two weeks of anti-VEGF-A treatment did not change the levels of soluble Aβ40, non-soluble Aβ40, soluble Aβ42, or non-soluble Aβ42 in APP/PS1 mice (Supplementary Figure 3).

**Figure 3.**
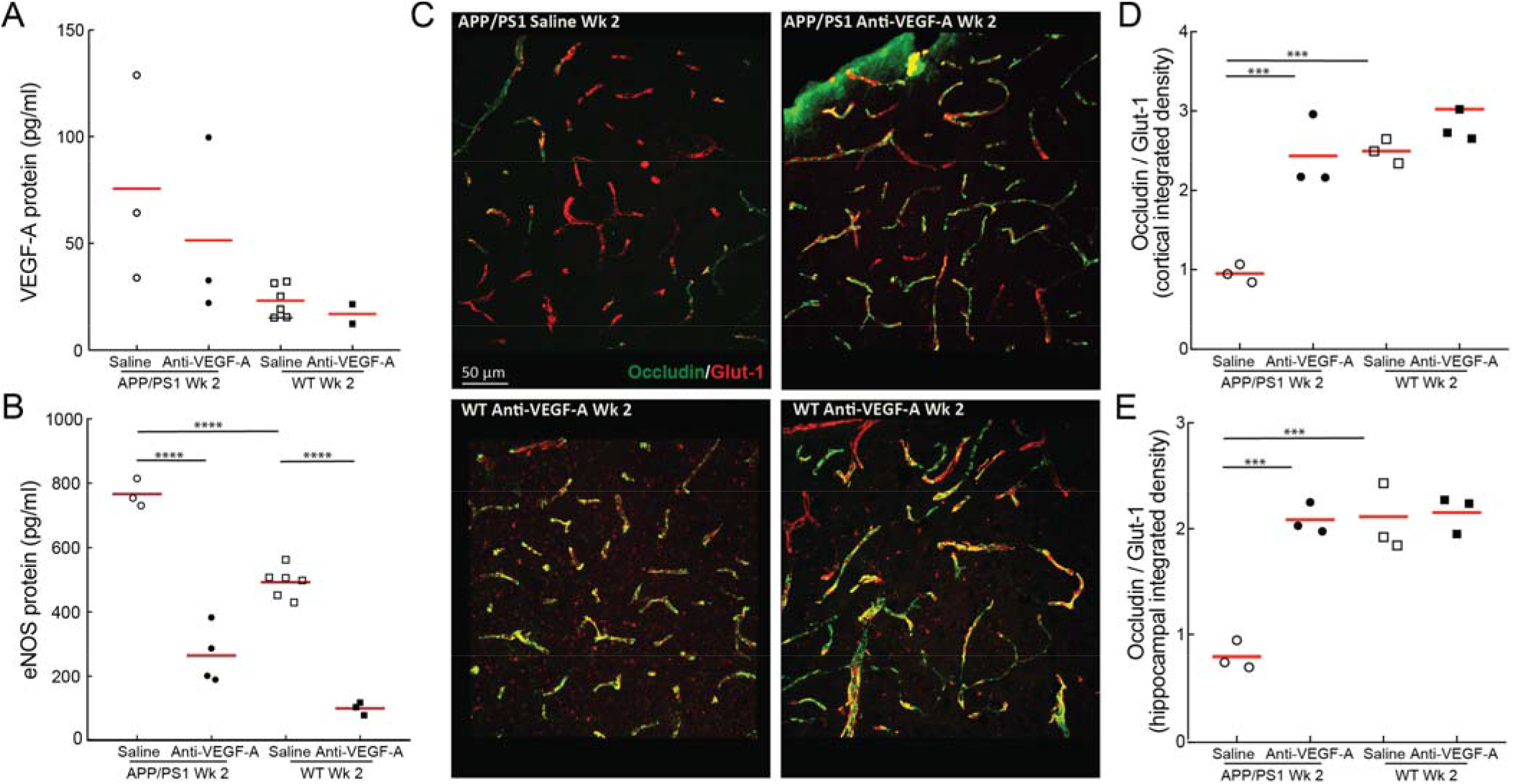
Anti-VEGF-A treatment modulates endothelial protein expression in APP/PS1 mice. ELISA measurements of VEGF-A (**A**) and eNOS (**B**) concentrations after 2 weeks of anti-VEGF-A treatment or saline control injections in APP/PS1 and WT mice (APP/PS1-saline: n=3, APP/PS1-anti-VEGF-A: n=4, WT-saline: n=6, WT-anti-VEGF-A: n=3; one way ANOVA with Tukey’s *post hoc* multiple comparison correction to compare across groups: APP/PS1-saline [VEGF-A] vs. APP/PS1-anti-VEGF-A [VEGF-A] p value < 0.001, WT-saline [VEGF-A] vs. WT-anti-VEGF-A [VEGF-A] p value = 0.81, APP/PS1-saline [VEGF-A] vs. WT-saline [VEGF-A] p value < 0.001, APP/PS1-anti-VEGF-A [VEGF-A] vs. WT-saline [VEGF-A] p value = 0.93, APP/PS1-saline [eNOS] vs. APP/PS1-anti-VEGF-A [eNOS] p value < 0.0001, WT-saline [eNOS] vs. WT-anti-VEGF-A [eNOS] p value < 0.0001, APP/PS1-saline [eNOS] vs. WT-saline [eNOS] p value < 0.0001). (**C**) Z-projection of confocal microscopy image stacks from representative cortical areas from mice of all four groups, revealing increased occludin density in anti-VEGF-A treated APP/PS1 mice as compared to saline injected APP/PS1 mice. Integrated density of occludin fluorescence as a function of the integrated density of the endothelial cell marker Glut-1 in the cortex (**C**) and hippocampus (**D**) (APP/PS1-saline: n=3, APP/PS1-anti-VEGF-A: n=3, WT-saline: n=3, WT-anti-VEGF-A: n=3; one way ANOVA with Tukey’s post hoc multiple comparison correction to compare across groups: APP/PS1-saline cortex vs. APP/PS1-anti-VEGF-A cortex p value = 0.0006, saline WT cortex vs. WT-anti-VEGF-A cortex p value = 0.53, APP/PS1-saline cortex vs. WT-saline cortex p value = 0.0005, APP/PS1-anti-VEGF-A cortex vs. WT-saline cortex p value > 0.99, APP/PS1-saline hippocampus vs. APP/PS1-anti-VEGF-A hippocampus p value = 0.0004, WT-saline hippocampus vs. WT-anti-VEGF-A hippocampus p value > 0.99, APP/PS1-saline hippocampus vs. WT-saline hippocampus p value = 0.0002, APP/PS1-anti-VEGF-A hippocampus vs. WT-saline hippocampus p value > 0.99). Each point represents one mouse and the red horizontal represents the median.

### Consistently stalled capillaries showed lower occludin expression in anti-VEGF-A treated APP/PS1 mice

To further explore the potential link between occludin expression and capillary stalling, we sought to correlate these on a capillary-by-capillary basis. In the anti-VEGF-A treated APP/PS1 mice, there were some capillaries that we observed to stall in the last imaging session (Figure 4A). In a total of five mice we were able to spatially align sagittal brain sections with the in vivo imaging by manually matching the patterns of the capillaries imaged in vivo with those labeled by post-mortem occludin staining (Figure 4B). Using this approach, we were able to unambiguously identify ten capillaries that were stalled on the last imaging session in the occludin stained sections. We further identified 100 capillaries that were flowing. We quantified the average intensity of occludin staining along identified capillaries and found that the stalled capillaries had ~40% lower occludin expression than the flowing capillaries in both anti-VEGF and saline treated APP/PS1 mice (Figure 4C) (Mann Whitney anti-VEGF p-value < 0.0001, saline p-value<0.001). The very low number of stalled capillaries in the anti-VEGF-A treated APP/PS1 mice coupled with the experimental challenge of unambiguously matching individual capillaries across the in vivo and post-mortem images precluded us from increasing the number of stalled capillaries that were analyzed.

**Figure 4.**
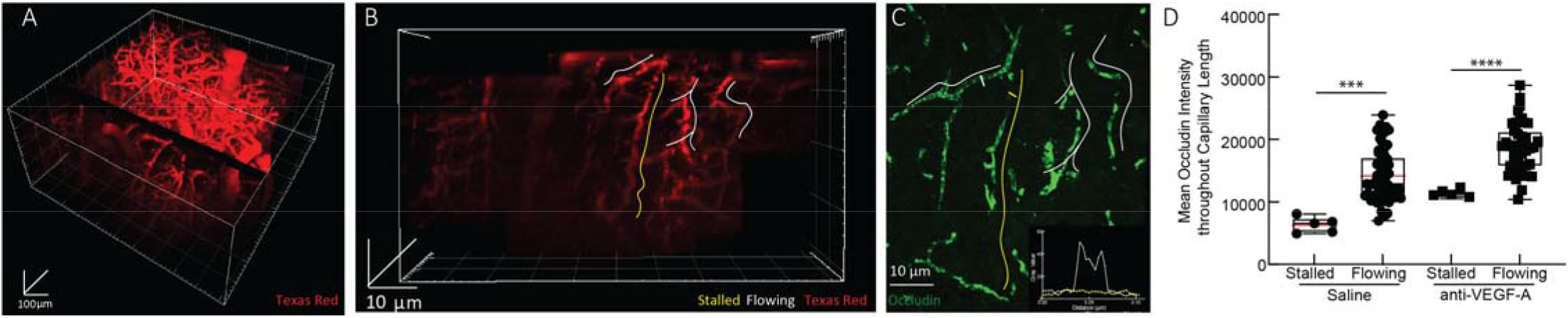
Anti-VEGF-A treated APP/PS1 mice had lower occludin expression in stalled capillaries as compared to flowing capillaries. Capillaries of APP/PS1 mice treated with anti-VEGF-A seen via *in vivo* 2PEF imaging (**A** and **B**), with flowing (white highlight) and stalled (yellow highlight) vessels indicated, spatially aligned to occludin immunofluorescence histopathology **(C)** revealing lower occludin fluorescence at stalled capillaries as compared to flowing capillaries. The inset graph in the lower right tracks the cross sectional mean grey value of occludin fluorescence across the distance of the white and yellow dashes on the stalled and flowing capillary. (**D)** Mean occludin concentrations, determined from the immunohistochemistry, for flowing and capillary segments, determined from *in vivo* imaging, in APP/PS1 mice treated with anti-VEGF-A for two weeks (APP/PS1 saline n=3 mice, 61 capillaries and APP/PS1 anti-VEGF-A n=2 mice, 43 capillaries; Kruskal-Wallis test with multiple comparison correction to compare across groups; saline p-value < 0.001, anti-VEGF-A p-value < 0.0001); bar graph represents mean values, and error bars represent standard deviation).

### Anti-VEGF-A treatment reduced capillary stalls and increased capillary blood flow speed in APP/PS1 mice within an hour of injection

VEGF-A signaling can rapidly change the incorporation of tight junction proteins, such as occludin, at the BBB on a ~15-minute time scale, altering BBB permeability (*36*). We tested whether the impact of anti-VEGF-A on the incidence of capillary stalls in APP/PS1 mice was similarly fast (Figure 5A). We found that one hour after a single anti-VEGF-A injection, APP/PS1 mice had 59% fewer capillary stalls (0.5%) than at baseline (1.2%) (repeated measures one-way ANOVA with Tukey’s *post hoc* multiple comparison correction; p value < 0.02) (Figure 5B - D). We continued giving anti-VEGF-A every other day for a week, and found that stall incidence remained low, consistent with the data in Figure 2. WT mice had low stall incidence both before and after anti-VEGF-A treatment (Figure 5D). At baseline, APP/PS1 mice had 25% slower capillary flow speeds, as measured with line scans (Figure 5E), as compared to WT mice. The rapid reduction in capillary stalls in APP/PS1 mice with anti-VEGF-A treatment was associated with a 20% on average, increase in capillary blood flow speed (vessels < 10 μm), at one hour after injection and was maintained at one week (Figure 5F). We did not observe differences in capillary diameter between groups nor did we detect significant changes in capillary diameter with anti-VEGF-A treatment (Figure 5G).

**Figure 5.**
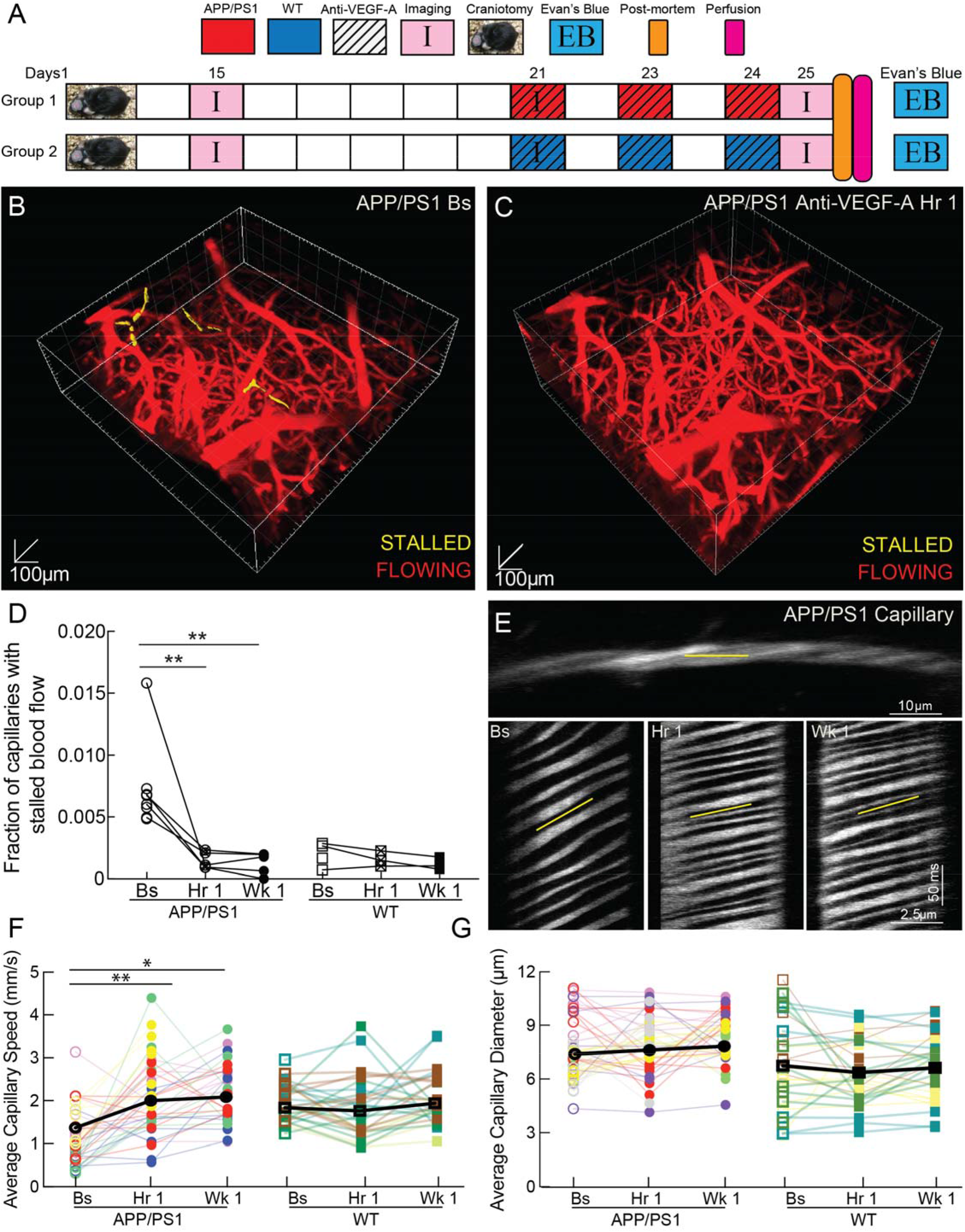
Anti-VEGF-A antibody treatment reduced capillary stalls and increased capillary flow speed within 1 hour in APP/PS1 mice. (**A**) Schematic of experimental timeline. Craniotomies were performed on APP/PS1 mice and they recovered for two weeks. The mice were divided into two groups: APP/PS1 mice treated with anti-VEGF-A (n=9; APP/PS1-anti-VEGF-A) and WT mice treated with anti-VEGF-A (n=4; WT-anti-VEGF-A). These mice were imaged three times – twice on the first imaging day before and ~1 hour after treatment and again five days later after two additional treatments. The brains were harvested for post-mortem assays after the second imaging session. Rendering of 2PEF stacks of the brain from the same APP/PS1 mouse before (**B**) and 1 hour after (**C**) anti-VEGF-A injection. Capillaries that were stalled are highlighted in yellow. (**D**) The fraction of capillaries with stalled blood flow at baseline (Bs), 1 hour after anti-VEGF-A injection, and after 1 week of treatment in APP/PS1 and WT mice ((APP/PS1: 4 mice could not be imaged at 1 week due to window loss or death); ~ 60,000 capillaries; repeated measures one-way ANOVA with Tukey’s post hoc multiple comparison correction to compare baseline to 1 hour and 1 week: APP/PS1-baseline vs. APP/PS1-1-hour p value = 0.02, APP/PS1-baseline vs. APP/PS1-1-week p value = 0.005; Kruskal-Wallis test with multiple comparison correction to compare across groups; each data point in the graph represents the capillaries with stalled blood flow as a function of total capillaries in 4-6 2PEF stacks for each mouse) (**E**) Image (top) and line-scans (below) from a representative capillary from an APP/PS1 mouse taken at baseline and at 1 hour and 1 week after anti-VEGF-A treatment, showing higher RBC flow speeds after treatment. (**F**) Average RBC flow speed and (**G**) vessel diameter from cortical capillaries at baseline and at 1 hour and 1 week after anti-VEGF-A injection in APP/PS1 and WT mice (APP/PS1-baseline: n=9, 53 capillaries, APP/PS1-1-hour: n=6 (1 mouse died, 2 mice lost cranial windows), 45 capillaries, APP/PS1-1-week: n=5 (1 mouse lost cranial window), 35 capillaries, WT-baseline: n=4, 21 capillaries, WT-1-hour: n=3 (1 mouse lost cranial window), 23 capillaries, WT-1-week: n=3, 21 capillaries; repeated measures one-way ANOVA with Tukey’s post hoc multiple comparison correction to compare baseline to 1 hour to 1 week: APP/PS1-baseline vs. APP/PS1-1-hour p value = 0.0100, APP/PS1-baseline vs. APP/PS1-1-week p value = 0.0288, WT-baseline vs. WT-1-hour p value = 0.9390, WT-baseline vs. WT-1-week p value = 0.2884). In all graph’s lines connecting data points represent the same capillary, while different colors represent individual mice.

### Anti-VEGF-A treatment reduced BBB permeability in APP/PS1 mice within an hour of injection

Consistent with the known BBB tightening effects of anti-VEGF-A treatment, we observed reduced staining of the brain with intravascularly circulating Evans Blue dye when these animals were sacrificed (Figure 6A). In summary, anti-VEGF-A treatment reduced capillary stalls and increased capillary blood flow speeds in APP/PS1 mice within one hour, bringing both to the levels of WT mice. We measured BBB permeability by quantifying the amount of intravenously circulating Evans Blue dye found in post-mortem, perfused brain tissue (Figure 6B). APP/PS1 mice treated with anti-VEGF-A one hour before injecting Evans Blue had half of the dye concentration in the cortex and hippocampus as compared to APP/PS1 mice treated with saline (One-way ANOVA with Tukey’s *post hoc* multiple comparison correction; p value < 0.01) (Figure 6C and D, respectively), which suggests that anti-VEGF treatment improved the BBB permeability of APP/PS1 mice to a similar level to that of WT mice.

**Figure 6.**
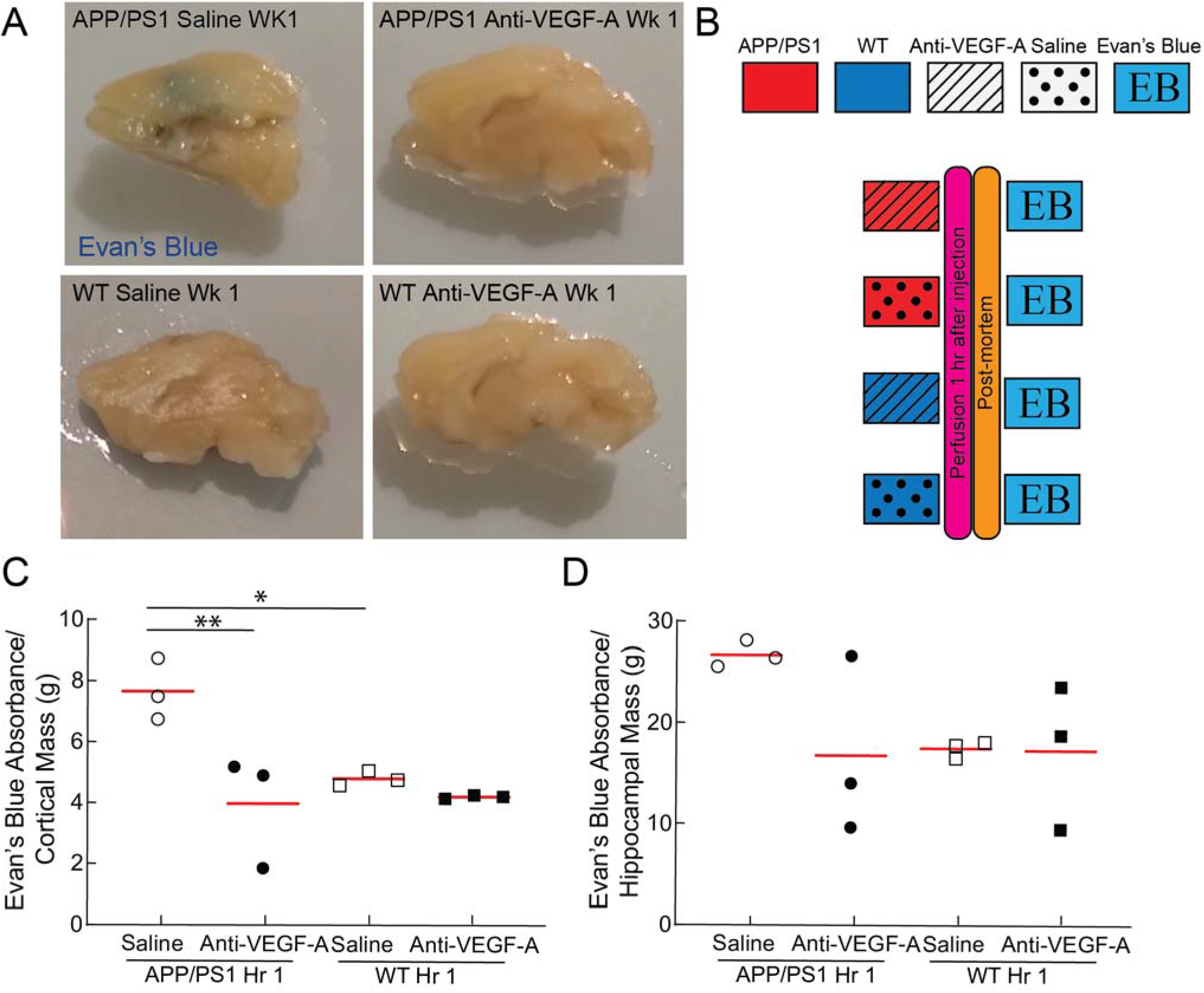
Anti-VEGF-A antibody treatment reduces BBB leakage within 1hr. in APP/PS1 mouse. (**A**) Sagittally cut brain slices from APP/PS1 and WT mice treated with anti-VEGF-A or saline for 1 week, and after 1 hour of intravenous circulation of Evan’s Blue dye. Arrow indicates region of increased penetration of Evan’s Blue into the brain in the APP/PS1, saline treated mouse. (**B**) Schematic of experimental timeline. APP/PS1 and WT mice were treated with anti-VEGF-A or saline. Evan’s Blue was intravenously injected an hour before perfusion and the extent of Evan’s Blue entry into the brain from the vasculature was quantified (n = 3 mice per group). Normalized absorbance of whole cortex (**C**) or hippocampus (**D**) tissue homogenates at 620 nm, to quantify the extent of Evan’s Blue entry into the brain for APP/PS1 and WT mice were treated with anti-VEGF-A or saline (one-way ANOVA with Tukey’s post hoc multiple comparison correction: panel C, saline APP/PS1 vs. anti-VEGF-A APP/PS1 p = 0.01; panel C, saline APP/PS1 vs. saline WT p = 0.04). In the graphs each point represents data from one mouse, and the red horizontal line represents the mean.

### Anti-VEGF-A treatment reduced capillary density in APP/PS1 mice after a week of chronic treatment

Finally, we probed whether the anti-VEGF-A treatment was associated with differences in cortical capillary density. We used DeepVess, a convolutional neural network-based segmentation algorithm, to segment 3D image stacks of the cortical microvasculature (excluding larger surface blood vessels) and quantified the fraction of image voxels that were within vessels to measure capillary density (Supplementary Figure 4A, B, C). APP/PS1 mice had a 20% higher capillary density than WT mice at baseline. One week of anti-VEGF-A treatment reduced capillary density by 8% in the APP/PS1 mice, bringing their capillary density in line with WT mice. Anti-VEGF-A treatment was not associated with capillary density differences in WT mice (Supplementary Figure 4C).

### Occludin level was reduced in patients with Alzheimer’s disease

Immunofluorescence of cortical tissue from AD patients and healthy controls demonstrated decreased occludin expression in the AD brain microvasculature (Figure 7A and B; Supplementary Table 1), similar to what we found in APP/PS1 mice.

**Figure 7.**
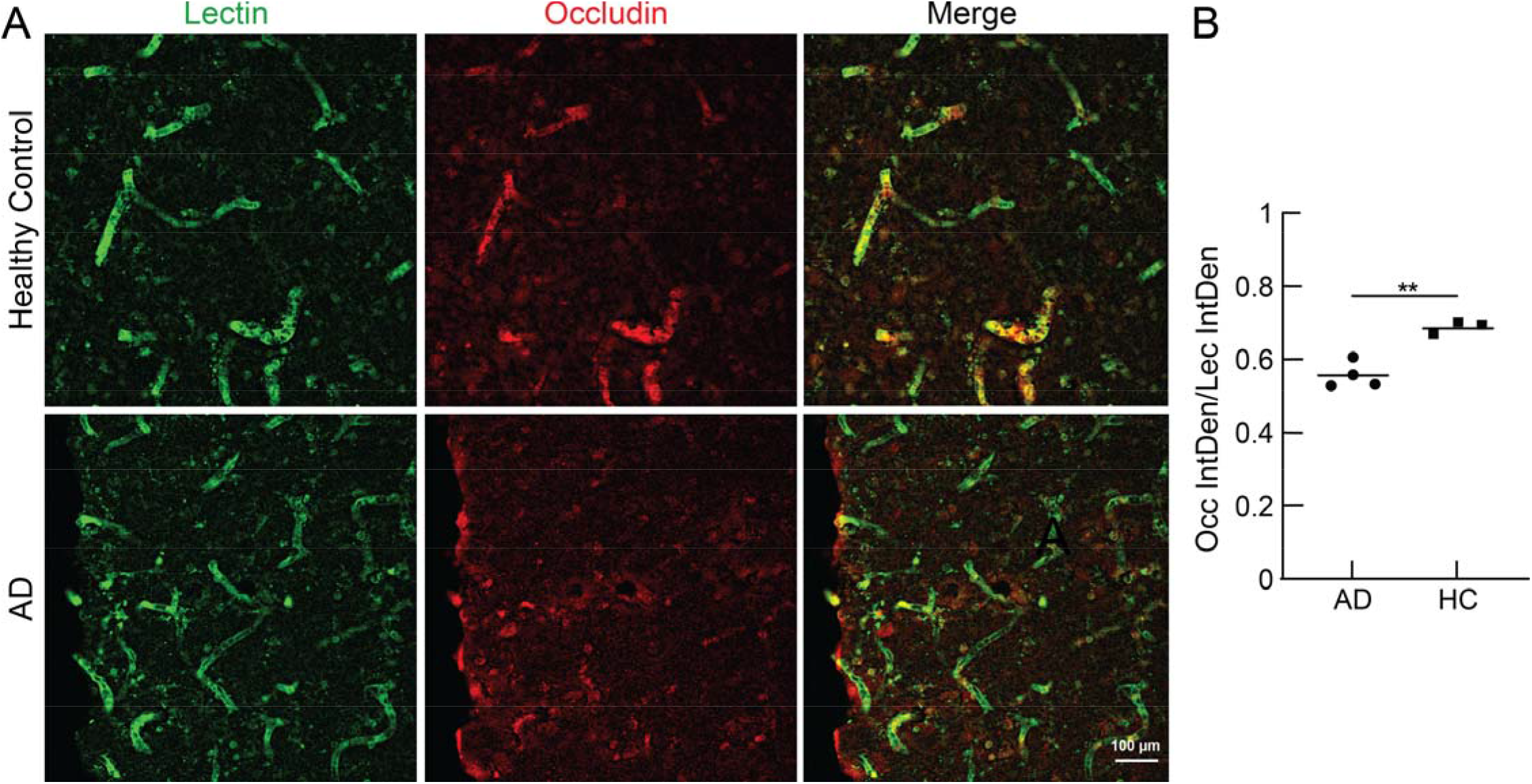
Occludin staining in cortical brain sections from AD patients and healthy controls. (**A**) Z-projection of confocal microscopy image stacks from representative cortical areas from healthy controls (upper panel) and AD patients (lower panel), revealing reduced occludin levels (red) in cortical capillaries identified by lectin staining (green). (**B**) Integrated density of occludin fluorescence, normalized to that of lectin. Eight fields of view from four sections were averaged for each sample. (Healthy controls n=3 and AD patients n=4; two-tailed unpaired t-test, ** indicates p = 0.01; red horizontal line represents the median)

## Discussion

CBF reductions occur in most dementia patients and are associated with cognitive impairment in cross-sectional studies (*37*). Such CBF reductions develop early in the pathogenesis of AD, when intervention would be most ideal (*38*). We previously showed that a small fraction of cortical capillaries is temporarily stalled at a four times higher rate in multiple mouse models of APP overexpression, as compared to WT mice (*37, 39, 40*). Furthermore, we found that administering an antibody against a neutrophil specific protein, Ly6G, led to a ~64% reduction in the incidence of capillary stalling and a ~17% increase in CBF. This was associated with rapid improvements in short-term memory in APP/PS1 mice as old as 16 months of age (*6, 41*). In this study we demonstrate that inhibition of VEGF-A signaling contributes to aberrant eNOS/occludin-associated BBB permeability, increases in capillary stalls, and reductions in CBF (Supplementary Figure 5).

A recent study followed the fate of capillaries with long-lived (> 20 min.) spontaneous or injected-microsphere-induced obstructions in WT mice, and found that obstructed capillaries were prone to being pruned from the vascular network, but that both obstructions and pruning were markedly decreased when VEGF-receptor 2 (the dominant VEGF-A receptor in endothelial cells (*42*)) signaling was blocked (*16*). Several studies have explored the role of VEGF-A signaling in patients with AD. Most studies find increased levels of VEGF-A in the serum or CSF of patients with AD (*18, 43–45*), with a minority showing reduced (*22, 46*) or unchanged levels (*47*). Increased VEGF-A levels are also seen in brain homogenates from the TgCNDR8 (*48*) and APP/PS1 mouse models of AD (*49*), as well as in endothelial cells of arteries, veins, and capillaries from aged WT mice(*50*), mouse models of APP over expression (*24, 51*), mutant Tau mice (*52*), and patients with AD (*53–55*). Our work here helps to tie these alterations in VEGF signaling in AD to capillary stalling and CBF reductions through experiments in APP/PS1 mice.

We showed that a two-week treatment of intraperitoneally injected anti-VEGF-A reduced the number of capillary stalls and increased capillary RBC flow speeds in APP/PS1 mice and that this was accompanied by a tightening of the BBB. Several cellular mechanisms are known to link VEGF signaling to BBB permeability, and these mechanisms may play a role in the inhibition of capillary stalling with anti-VEGF treatment. Luminal VEGF-A signaling up-regulates enzymes, including the endothelial isoform of nitric oxide synthase (eNOS), that increase vascular permeability, allowing for the endothelial cell rearrangements that support sprouting angiogenesis (*56*). The eNOS upregulation leads to decreased expression and modifications of several tight junction (TJ) proteins, including occludin (*57*). Decreased occludin levels are associated with increased BBB permeability (*58, 59*) and decreased TJ complex pore formation (*60*). In addition, eNOS knockout mice show increased levels of TJ proteins including occludin, and have decreased BBB permeability (*61*). Here, we found increased eNOS and decreased occludin expression in APP/PS1 mice, which was rescued with anti-VEGF treatment. Moreover, in a select dataset that correlated in vivo stalls with post-mortem protein expression, we also showed that occludin levels are reduced specifically in stalled capillaries, further strengthening this correlation. This finding is consistent with previous results showing decreased occludin expression in AD mouse models, including the APP/PS1 model (*23, 62*), and in the cortex of AD patients (*63, 64*), which we replicated here. We also observed very fast effects on capillary stalling and CBF with anti-VEGF-A treatment in APP/PS1 mice, with these rapid changes in capillary stalling and CBF of the same magnitude as those seen with 1-2 weeks of anti-VEGF-A administration. The rapid changes in capillary stalling were, again, correlated with decreased BBB permeability. Interestingly, increases in the phosphorylation status of occludin are implicated in decreased BBB permeability that occurs within minutes of inhibiting VEGF-A signaling (*36, 65, 66*). It seems likely that such post-translational modifications similarly contribute to the rapid changes in capillary stalling seen here. Other downstream effects of VEGF signaling, such as angiogenesis (*7*), vascularization (*8*), lymphangiogenesis (*9*), growth tip guidance (*10*), neurogenesis (*10*), and neuroprotection, are less likely occurring at this rapid timescale, further implicating loss of BBB integrity and hyperpermeability in the formation of capillary stalls. Indeed, changes in capillary density were only seen after 1 week of anti-VEGF therapy among APP/PS1 mice.

An analogous supporting these ideas has been suggested in the retina. Several studies have analyzed the role of VEGF-A signaling in the blood-retina barrier (BRB) permeability in diabetic retinopathy (DR) and have shown that intravitreal injections of anti-VEGF-A lead to an increase in occludin phosphorylation and tightening of the BRB within hours that was correlated with a reduction in the incidence of leukocytes stuck in retinal microvessels (*66–69*). In DR inhibition of the vascular inflammatory receptor ICAM-1 leads to reduced leukostasis and vascular leakage (*70*), and VEGF-A leads to increased levels of ICAM-1 in the vasculature (*71*). In AD low level chronic vascular inflammation is also present (*72*). This is supported by recent single-nucleus transcriptome analysis of endothelial cells, which have shown an increased level of genes involved in vascular inflammation in aged healthy humans (*50*), and patients with AD (*54*). The functional consequences of increased chronic vascular inflammation have also been studied. For example, mouse models of AD have increased endothelia/leucocyte interactions due to increased inflammation and the presence of neutrophil extracellular traps. (*73–75*). Ultimately, this chronic vascular inflammation could directly increase the number of capillary stalls and impact CBF. Overall this data indicates that luminal VEGF-A signaling plays a highly complex role in regulating endothelial function in the APP/PS1 mouse.

VEGF plays a role in a wide variety of signaling pathways and can act both on endothelial cells (luminal) and brain cells (parenchymal). Alterations of VEGF signaling in different tissue compartments of the brain and in the periphery could have different impacts on downstream aspects of AD, such as cognition (*76*). In this study, we measured increased levels of VEGF-A in APP/PS1 mice from brain lysates. Unfortunately, we could not separate our analysis into cell-types - endothelial, neuronal, or glial - due to a lack of raw material. This study has linked peripheral VEGF-A signaling to increased rates of capillary stalling and CBF reductions in the APP/PS1 mouse model of AD. However, VEGF-A is a multifactorial signaling molecule and has various functions in different cell-types. For example, VEGF-A supplementation led to improved short-term memory in mouse models of AD (*77*). The injection of bone marrow derived mesenchymal stem cells expressing the human VEGF-A165 isoform led to cognitive improvements in the plus-maze discriminative avoidance task, reduced parenchymal amyloid plaques, and initiated neovascularization (*78*). Similar results were seen in mice over-expressing the human VEGF-A165 in neurons in mouse models of AD (*77, 79*). Overall this sounds contradictory, but we show that reducing peripheral levels of VEGF-A increases BBB integrity and improves CBF, while others show over-expressed neuronal VEGF-A via stem cells transplantation leads to improvements of AD-related pathology. However, this could simply mean that increased VEGF-A signaling may be protective to the CNS on one side of the blood brain barrier (BBB) and that it may contribute to mechanisms within the vasculature that enhance the impact of AD pathology. Unifying these hypotheses suggests that increased levels of peripheral VEGF is a compensatory mechanism in response to reduced signaling in the brain. The idea that VEGF-A is still produced but the signaling is impaired has been proposed as pathological angiogenesis in AD (*80*). Furthermore, this hypothesis is also supported by the fact that large molecules, including IgG immunoglobulins (MW=150kDa), have a less than 1 in 1000 probability to cross the BBB (*26, 28*), even in BBB impaired models such as the APP/PS1 model (*81*). This suggests, that the anti-VEGF-A 165 antibody disproportionally inhibits luminal rather than parenchymal VEGF-A signaling.

In summary, this study describes how increased peripheral VEGF-A levels in the microvasculature of APP/PS1 mice lead to impaired BBB integrity, which correlates with increased capillary stalling, and reduced CBF. Furthermore, it describes how increased non-functional, pro-angiogenic VEGF-A signaling seen in AD could contribute to reduced CBF likely via capillary stalls.

## Acknowledgements

We want to thank Mike Lamont for developing the DeepVess editor app. Confocal imaging data was acquired through the Cornell University Biotechnology Resource Center, with NIH funding (grant number RR025502) for the shared Zeiss LSM 710 Confocal. Human brain tissue was provided by the NIH NeuroBioBank, the Harvard Brain Tissue Resource Center, Sepulveda Research Corporation, the Mount Sinai/JJ Peters VA Medical Center NIH Brain and Tissue Repository, and the University of Maryland Brain and Tissue Bank. This research was supported by the German National Academic Scholarship Foundation (KF), the National Institutes of Health grants AG049952 (CBS) and NS108472 (CBS), and the BrightFocus Foundation (CBS), and by the DFG German Research Foundation (OB).

## Supplementary Figures

**Supplementary Figure 1.**
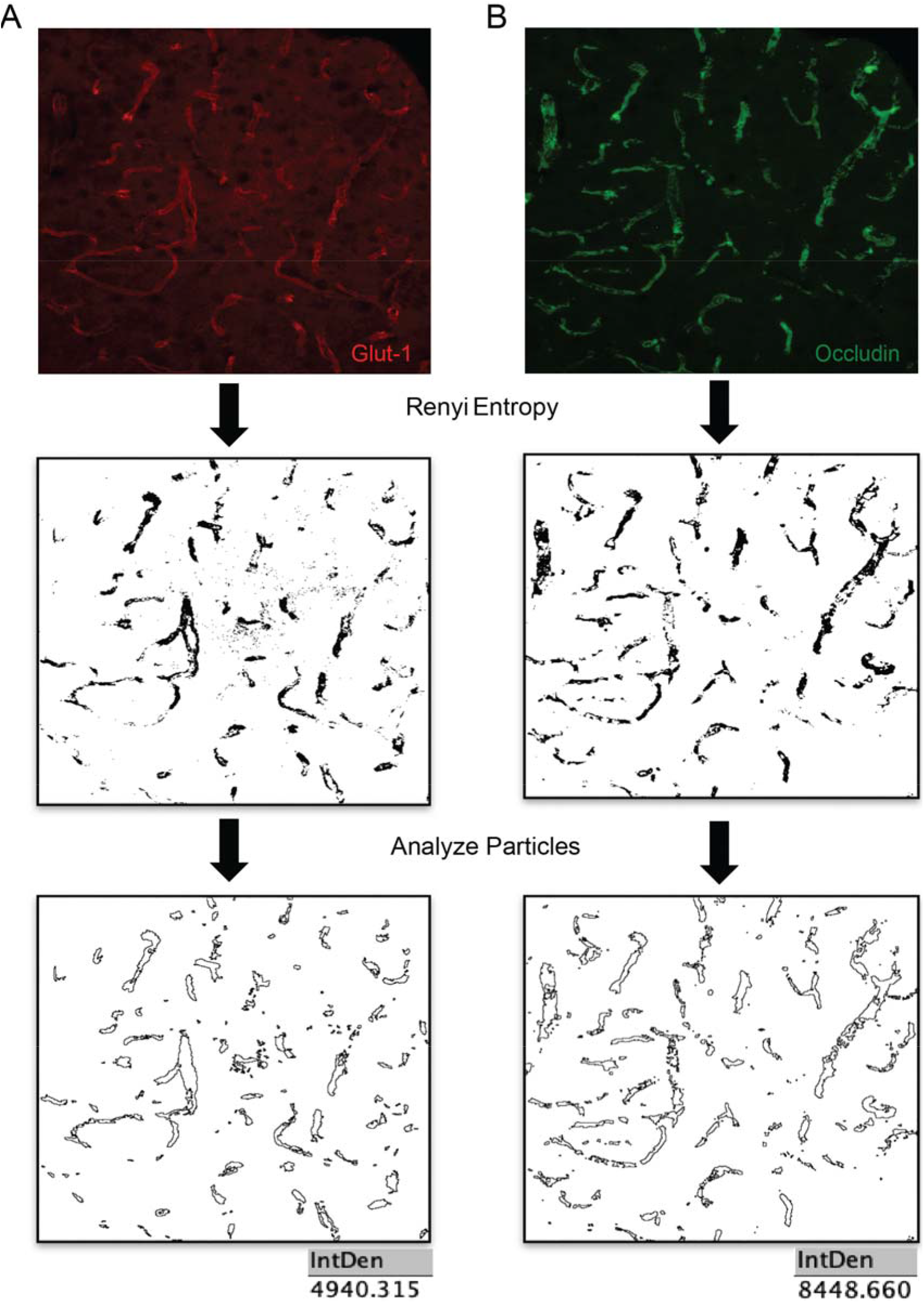
Renyi entropy filter and integrated density ratio was used to determine occludin concentrations at capillaries. Projections of confocal images stacks of occludin and glut-1 labeled cortical sections (top) were subjected to a Renyi entropy filter to separate background from antibody stained pixels (middle) and particles with radius larger than 5 pixels were counted (bottom) for the vascular marker glut-1 (**A**) and for occludin (**B**), and the ratio of these was computed.

**Supplementary Figure 2.**
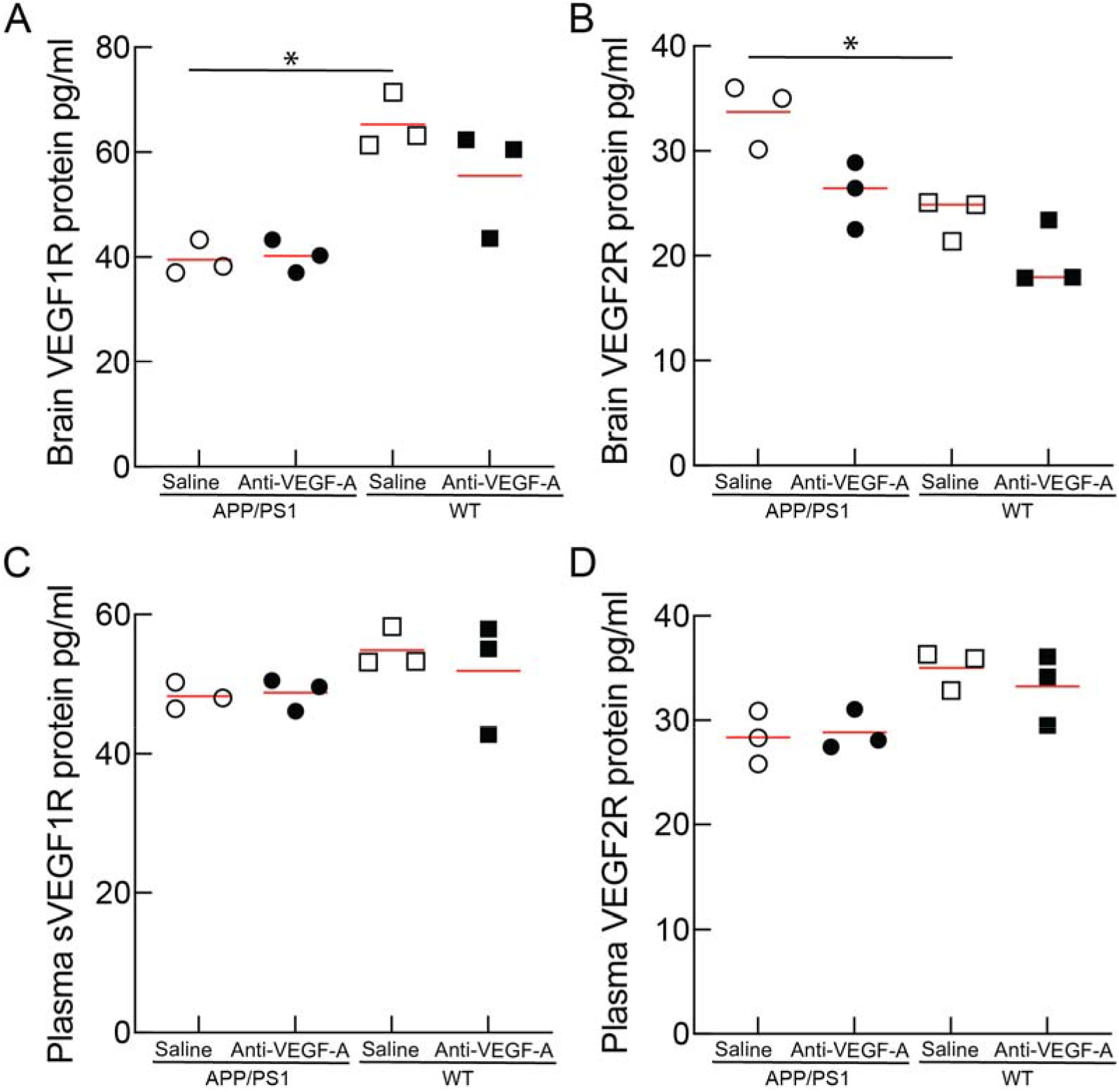
VEGF 1 and 2 receptor protein levels from brain and plasma samples after anti-VEGF-A antibody treatment. ELISA measurements of VEGF1R receptor (**A**), VEGF2R receptor (**B**) from brain lysates, and soluble VEGF1R receptor (**C**), and soluble VEGF2R receptor (**D**) from plasma samples after 1 week of anti-VEGF-A treatment or saline injections in APP/PS1 and WT mice (n=3 mice per group). Kruskal-Wallis test with multiple comparison correction to compare across groups. In all graphs each point represents the ELISA measurement from one mouse, red horizontal line represents the median; * indicates p = 0.05.

**Supplementary Figure 3.**
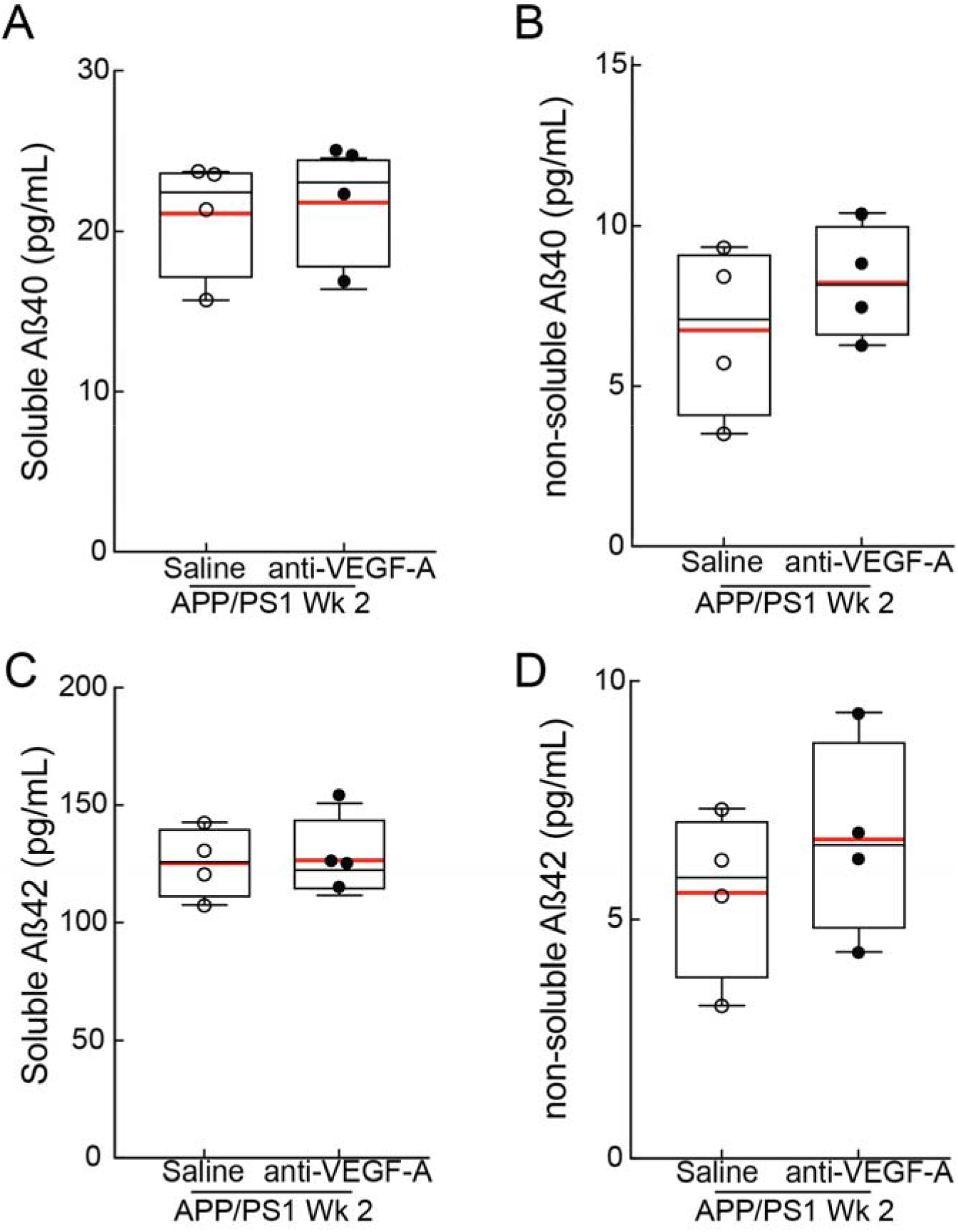
Amyloid-Beta (Aβ) levels were not modulated by anti-VEGF-A antibody treatment. ELISA measurements of soluble amyloid-Beta (Aβ) 40 (**A**), non-soluble Aβ40 (**B**), soluble Aβ42 (**C**), and non-soluble Aβ42 (**D**) concentrations after 2 weeks of anti-VEGF-A treatment or saline injections in APP/PS1 and WT mice (n=4 mice per group). In all graphs each point represents the ELISA measurement from one mouse, the boxplot whiskers extend 1.5x the difference between the 25^th^ and 75^th^ percentile of the data, the red horizontal line represents the median, and the black line represents the mean.

**Supplementary Figure 4.**
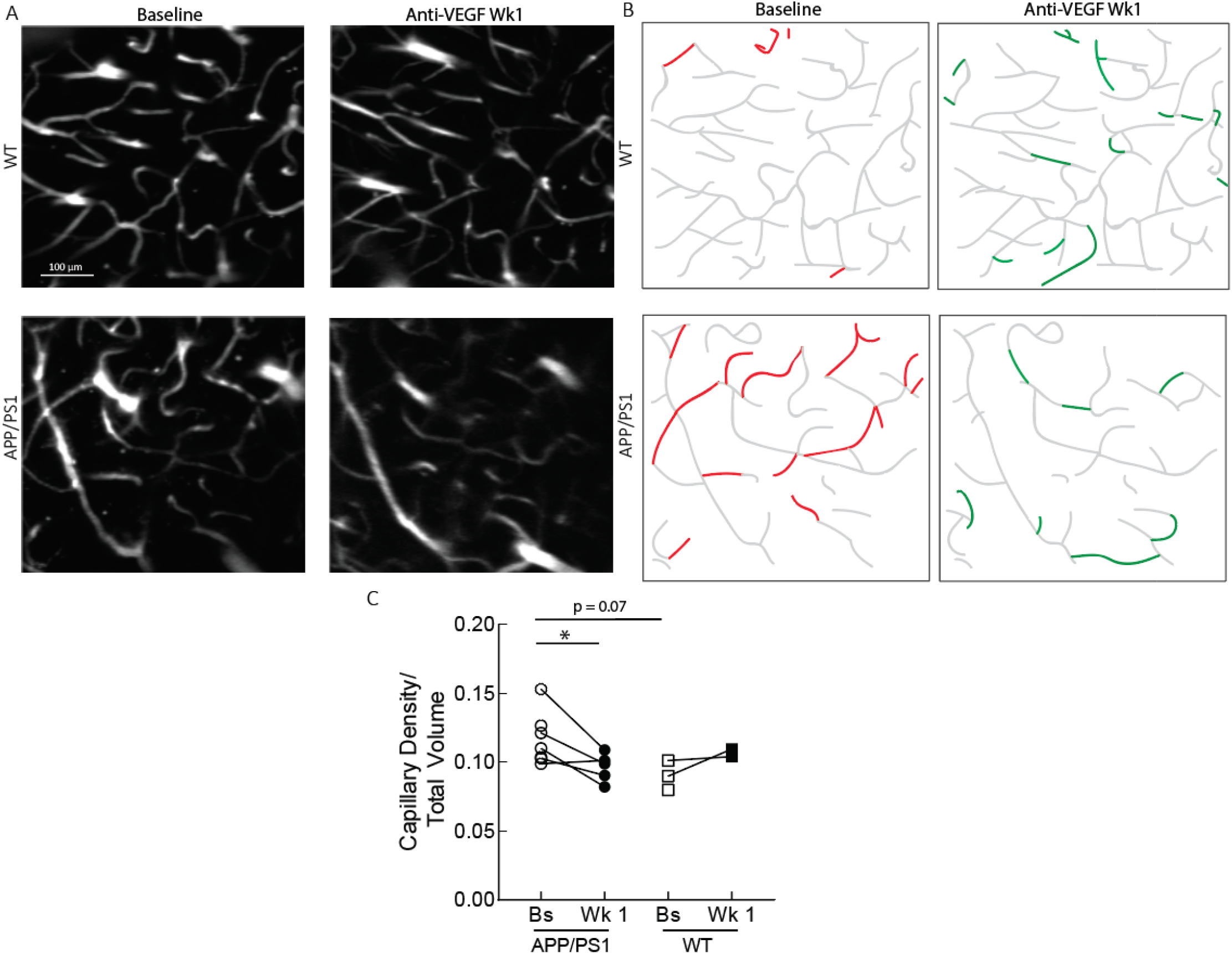
Anti-VEGF antibody treatment reduces capillary density after 1 week of treatment in APP/PS1 mice. **A** 40 μm deep z-projections from one APP/PS1 and one WT mouse at baseline and a week after chronic anti-VEGF treatment, showing the loss of multiple capillaries in the APP/PS1 mouse, but few in the WT mouse. **B.** Capillary traced to mark the loss (red) and gain (green) of capillary segments after 1 week of anti-VEGF treatment. **C** Capillary density in APP/PS1 and WT mice measured by automated segmentation before and after anti-VEGF treatment (baseline APP/PS1: n=7 (1 mouse’s window was not clear enough for 2PEF image stacks, 1 mouse was imaged at a slightly different zoom and was thus excluded from this analysis), 1 week APP/PS1: n=5 (1 mouse died, 1 mice lost cranial window), baseline WT: n=3 (1 mouse was imaged at a slightly different zoom and was thus excluded from this analysis), 1 week WT: n=2 (1 mouse lost cranial window); paired t-test to compare baseline to 1 week: baseline APP/PS1 vs. 1 week APP/PS1 p value = 0.0430, baseline WT vs. 1 week WT p value = 0.4051; one way ANOVA with Tukey’s post hoc multiple comparison correction to compare across groups: baseline APP/PS1 vs. baseline WT p value = 0.0715, 1 week APP/PS1 vs. baseline WT p value > 0.99; each point in the graph represents one mouse, with the line connecting them representing the same mouse tracked over a week).

**Supplementary Figure 5.**
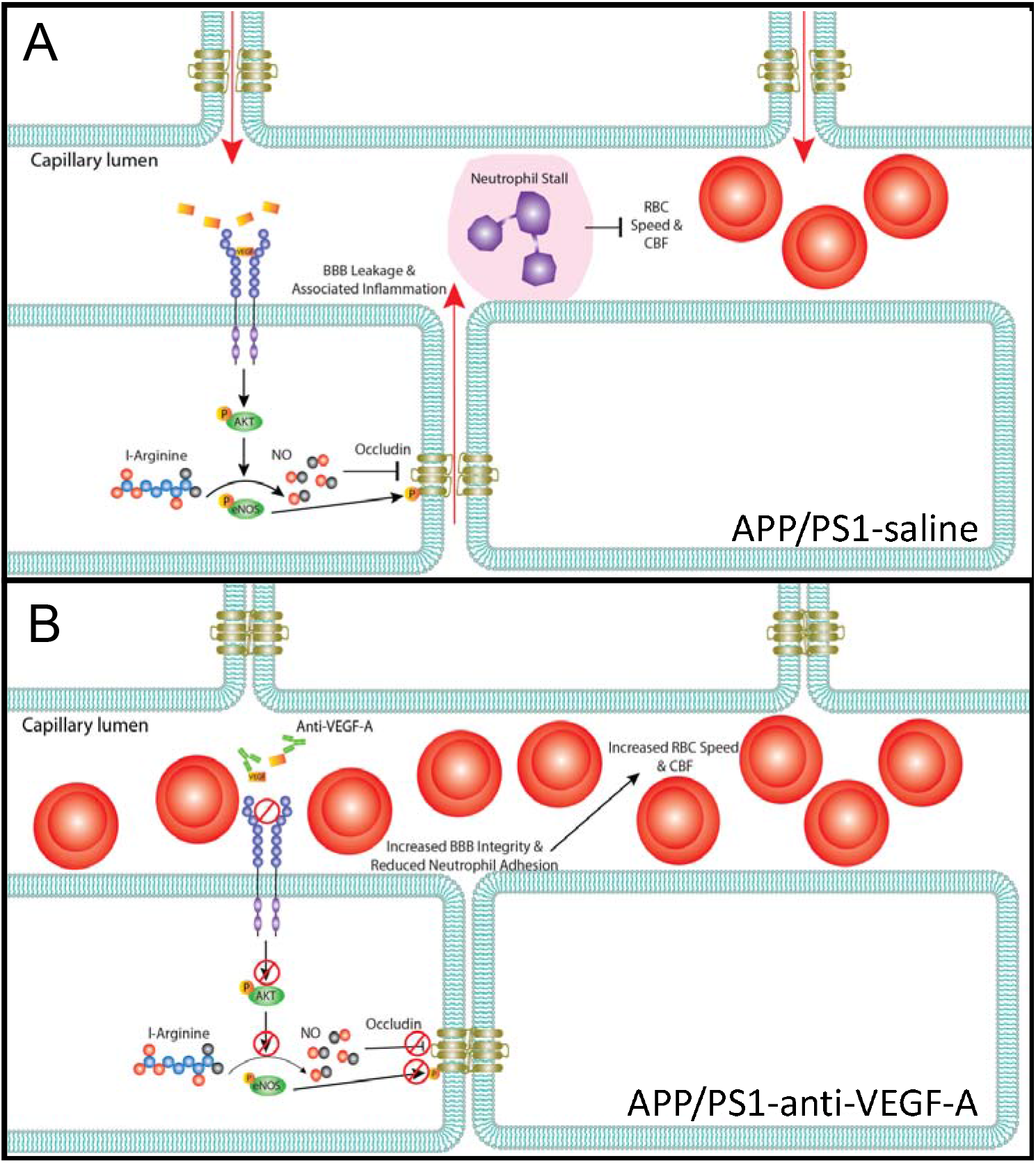
Proposed mechanism linking VEGF-A signaling and leukocyte stalling in APP/PS1 capillaries. **(A)** In APP/PS1 mice an increased fraction of capillaries are stalled. Pathological angiogenesis is attributed to AD. The hypothesis suggests that endothelial cells respond to increased amyloid-beta through increased ROS levels, causing vascular damage. Finally, this leads to an increase in angiogenic cytokines, such as thrombin and VEGF-A. Long-term VEGF-A exposure leads to BBB leaks (red arrows) and damages vessels in ways that may contribute to capillary stalling. (B) Here we show that peripheral treatment with an anti-VEGF-A antibody reduces the number of capillary stalls and increases CBF, which was associated with reduced VEGF-A levels and reduced BBB leakage.

**Supplementary Table 1.** Demographics of human brain tissue samples.

| **AGE** | **SEX** | **PHENOTYPE** | **BRAIN AREA** |
| --- | --- | --- | --- |
| 69 | M | Unaffected Control | Cortex: Middle temporal (Brodmann area 21) |
| 82 | F | Alzheimer's disease with late onset | Cortex: Middle temporal (Brodmann area 21) |
| 75 | F | Alzheimer's disease, unspecified | Cortex: Middle temporal (Brodmann area 21) |
| 79 | M | Unaffected Control | Cortex: Middle temporal (Brodmann area 21) |
| 70 | F | Unaffected Control | Cortex: Middle temporal (Brodmann area 21) |
| 80 | F | Alzheimer's disease, unspecified | Cortex: Middle temporal (Brodmann area 21) |
| 84 | F | Unaffected Control | Cortex: Middle temporal (Brodmann area 21) |
| 62 | M | Alzheimer's disease, unspecified | Cortex: Middle temporal (Brodmann area 21) |
| 89 | M | Alzheimer's disease, unspecified | Cortex: Middle temporal (Brodmann area 21) |

